# Prescribed fire selects for a pyrophilous soil subcommunity in a northern California mixed conifer forest

**DOI:** 10.1101/2022.04.15.488482

**Authors:** Monika S. Fischer, Neem J. Patel, Phillip J. de Lorimier, Matthew F. Traxler

## Abstract

Low intensity prescribed fire is a critical strategy for mitigating the effects of catastrophic wildfires. The above-ground response to fire has been well-documented, including many ecosystem benefits associated with prescribed burning, but fewer studies have directly addressed the effect of prescribed fire on soil organisms. We aimed to understand how soil microbial communities respond to prescribed fire and to determine the ecological processes driving their dynamics. We extensively sampled four plots for 17 months in a mixed conifer forest in northern California, USA; immediately following a low-intensity prescribed fire, a higher-intensity prescribed fire, and two no-burn control plots. We found that prescribed fire significantly altered the community structure for both fungi (ITS) and bacteria (16S), which was sustained throughout the time-series. By comparing our community profiling results with a model of neutral community assembly, we found that the presence of most taxa across all experimental conditions could be explained by neutral processes. However, combining threshold indicator taxa analysis and correlation network analysis with the neutral model identified a cohort of taxa that responded deterministically to prescribed fire. The subcommunity identified through this series of analyses includes both known and new pyrophilous taxa. Beyond this, our analyses revealed network modules within postfire communities which were responsive to fire-intensity. Taken together, these results lay the foundation for building a process-driven understanding of microbial community assembly in the context of the classical disturbance regime of fire.

## INTRODUCTION

Under the current trajectory of climate change, wildfires are expected to continue increasing in frequency and severity in western North America and other regions across the globe [1–3]. Wildfires burn hundreds of millions of hectares of vegetation annually [4] and contribute to the global carbon cycle both in terms of atmospheric input as CO_2_ and terrestrial sequestration of carbon in the form of pyrogenic organic matter (PyOM) [5–7]. A key strategy toward decreasing the frequency of devastating wildfires is the implementation of lower-intensity prescribed fires and managed wildfires. These lower-intensity controlled burns have proven effective in reducing the fuel load and the likelihood of catastrophic fire [8–10]. While the effects of prescribed fire on plant and animal communities are increasingly well-documented [9–13], our understanding of the impact of fire management strategies on microbial communities is much more limited [14]. In this study, we investigated the effects of prescribed burns on soil bacteria and fungi with the aim of understanding the ecological processes that shape these communities.

Several key ecological processes operate together to dictate community assembly after perturbations such as fire. One process by which communities assemble is the deterministic (or ‘niche’) process of ecological selection, which encompasses environmental filtering and biotic competition. In contrast, neutral (or random) processes of community assembly include dispersal, drift, and speciation [15]. Understanding the relative contributions of these processes to community assembly after disturbance is critical because; (1) they ultimately determine the successional trajectory and recovery time of these communities, and (2) possible interventions designed to enhance community resilience will likely need to take these processes into account if they are to be successful. When a new habitat becomes available for microbial colonization, neutral processes typically dominate early on, with selection playing a more important role over time [16, 17]. Fire can be among the most extreme types of disturbance, effectively resulting in a new habitat that is recolonized over time via secondary succession [18]. Consistent with this notion, Ferrenberg, *et al* found that neutral processes played a strong role in soil bacterial communities four weeks after severe wildfire, and after sixteen weeks the community was less neutral [17].

Work across multiple post-fire systems (predominantly wildfire) is starting to paint a complex picture of the impacts of fire on microbial communities. Despite complete decimation of the above-ground ecosystem, many organisms are able to survive below ground, even during the most extreme wildfires [12]. A recently proposed conceptual model provides a framework for considering the thermo-chemical gradient that defines the post-fire soil habitat [19]. This model describes the insulative capacity of soil, meaning that the effect of fire decreases with depth, and organisms below a certain depth are protected from the heat of the fire by the soil itself. Despite survival at deeper depths, the organisms present near the soil surface are generally dramatically impacted by fire. The documented effects of fire on the soil microbial community include a decrease in overall biomass [20–22] and a significant perturbation of the community structure [23–27]. Both fungal and bacterial communities have been observed to decrease in richness after severe wildfire [23–25, 27], although there are notable exceptions [26].

Over a hundred years of macroscopic observation [28–30], combined with recent community DNA sequencing studies, suggest that a distinct pyrophilous fungal community assembles in wildfire-affected soils. These pyrophilous fungi produce abundant fruiting bodies commonly seen on the surfaces of burnt soil and pyrolyzed wood, and more recently these taxa have also been identified via DNA sequencing of post-fire soils [19, 24–27, 31–38]. Commonly observed pyrophilous fungi include members of the genera *Pyronema, Anthracobia, Geopyxis, Tricharina, Morchella, Peziza, Pholiota, Lyophyllum, Myxomphalia*, and *Neurospora*. Comparatively less is known about bacterial taxa that respond positively in post-fire soils, although taxa that appear across recent studies include *Paraburkholderia, Arthrobacter, Flavobacterium*, and *Massilia* [25–27]. To our knowledge, only one recent study has investigated the effect of prescribed fire on both soil fungal and bacterial communities. Mino *et al* conducted ten replicate low-and high-intensity prescribed fires in a shrub-encroached prairie ecosystem, and analyzed samples from two time points; pre-fire and post-fire [31]. They found that high-intensity fire, but not low-intensity fire, reduced richness of both bacteria and fungi. Additionally, the bacteria *Massilia, Domibacillus*, and the fungi *Neurospora, Pyronema, Anthracobia*, and *Penicillium* were significant indicators of post-fire samples [31]. Beyond this example, soil microbial community assembly after prescribed burns has not been deeply explored.

In this work, we sought to address two major questions. First, does prescribed fire lead to assembly of a community of pyrophilous organisms similar to those seen after severe wildfires? And second, what ecological processes drive formation of the community after prescribed fire? To answer these questions, we deeply sampled two prescribed burn plots and two no-burn control plots over 17 months in a mixed conifer forest in the Sierra Nevada mountains of northern California, USA. Combining threshold indicator taxa analysis (TITAN), correlation network analysis, and a model of neutral community assembly delineated a group of taxa whose presence is likely explained by a deterministic response to prescribed fire. Importantly, many members of this pyrophilous subcommunity have been previously described as responding positively after wildfire, and our analyses add several new taxa to the ranks of potentially pyrophilous microbes. While this pyrophilous subcommunity likely assembled as the result of selective processes following prescribed fire, we find that the majority of the postfire community is likely assembled as the result of neutral processes. This work provides a foundation for building a mechanistic understanding of the ecological processes that shape post-fire microbial communities.

## MATERIALS & METHODS

### Prescribed fire treatments

Prescribed fires were conducted at the University of California’s Blodgett Forest Research Station located near Georgetown, CA, USA. Blodgett Forest is a mixed conifer forest (mostly *Pinus lambertiana, Pinus ponderosa*, and *Pseudotsuga menziesi*, with some scattered *Calocedrus decurrens, Sequoiadendron giganteum, Abies concolor*, and *Quercus kelloggii*). Soil is in the Holland series, generally characterized as fine-loamy, mixed, mesic Ultic Haploxeralfs. We established four 10m transects within the forest at roughly 1360m elevation; Hi (38.89598, −120.64800), Lo (38.90016, −120.65648), 1c (38.90562, −120.66345), and 2c (38.90191, −120.65901). The Hi plot was treated with a high-intensity slash-pile burn on 4 Jan 2019. The Lo plot was treated with a low-intensity broadcast burn on 25 Oct 2018, and plot 2c was treated with a moderate-intensity broadcast burn on 13 Feb 2020. All prescribed burns were facilitated with fossil fuel drip torches. Fossil fuel was excluded from transects, instead, ignited fossil fuel was dripped at least two meters away, and fire naturally traveled across our transects as it burned through dry plant debris. To measure soil temperature during and after fire, we buried Extech SDL200 data loggers ∼0.5m below the soil surface, and ∼2-3m away from our sampling transect. Each data logger was equipped with four thermocouples, and the data loggers were protected inside a hard plastic shoebox with a hole cut in the side to allow the thermocouples to exit. Temperature was measured every minute until the batteries died (roughly four days).

### Sample collection

Transect sampling locations were randomly assigned without replacement for each plot for the entirety of our sampling time-series. Prior to burning, and in control plots, 10cm soil cores were collected using an ethanol-sterilized soil sampler (JMC PN031). After burning, we collected soil core samples from 0-3cm and 3-6cm. Sampling depths were initially determined based on previous work [19] and the data from our thermocouples. We also conducted a pilot experiment in plot 2c to directly test if sampling depth affected the resulting observed microbial community. For this pilot experiment, six replicate soil samples were collected from the following depths at two time points pre- and post-fire; 0-1cm, 1-2cm, 2-3cm, 3-4cm, 4-5cm, 5-10cm, and 10-20cm. In addition, we later pooled equivalent amounts of soil from 0-3cm and from 0-10cm prior to DNA extraction. Figure S1 and S2 illustrate that there were no significant differences in the composition or diversity of the microbial communities observed at the scale of 0-20cm in Blodgett soil. (PERMANOVA p > 0.05, and ANOVA p > 0.1, n = 6).

All plots were sampled at every time point. Triplicate soil samples were collected at least once immediately prior to burning, once immediately after burning, and then once/month thereafter. To capture community dynamics associated with the first precipitation event, we increased our sampling frequency during the weeks following the first precipitation event and the start of the wet season in late November 2018. Multiple cores were collected per sample and immediately pooled in a 50mL conical tube, to a total volume of ∼25-30mL of soil. After collection, samples were transported by car and immediately placed in a −80 ºC freezer. pH was measured using a soil to water ratio of 1:2.5 within 48 hours of sample collection.

### DNA extraction, PCR amplification, and sequencing

We generally followed the methodology described by Simmons, *et al* [39]. To isolate total gDNA from soil, 1.2-1.5g of soil was transferred to a 2mL eppendorf tube and then we followed the Qiagen PowerSoil DNA extraction kit protocol. We eluted the gDNA on 100ul of DEPC water. 5ng/ul of DNA was used for simultaneous PCR amplification and illumina library prep with dual 12bp barcodes. Primer sequences in File S1 [39–41]. Samples were randomly assigned barcodes, and randomly distributed spatially across PCR plates (excluding corners) and sequencing libraries. 100ng of DNA from each sample was pooled to form a library. Libraries were sequenced via a PE300 strategy on illumina MiSeq, and then the resulting data were demultiplexed at the UC Davis Genome Center.

### Raw sequence processing

For the ITS sequences, we used AMPtk v1.5.1 (which used VSEARCH v2.15.0) to quality filter reads, trim off primer sequences, and merge forward and reverse reads together [42]. Merged reads less than 100bp were filtered out, and remaining reads were trimmed to 300bp. We then used DADA2 v1.14 (via R v4.0.3) to dereplicate the sequences, infer Amplicon Sequence Variants (ASVs), remove chimeras, assign taxonomy, and ultimately build the tables used in downstream analyses: OTU table, taxonomy table, and sequence table [43, 44]. The UNITE v.8.3 database was used to assign fungal taxonomy, and FUNGuild added functional guild information for many taxa [45, 46]. For 16S sequences, we used Quantitative Insights into Microbial Ecology version 2 (QIIME2 v2021.8) for all our read processing steps [47]. We used built-in tools to quality filtered reads and to trim reads where the quality score was < 25. QIIME2 incorporates cutadapt [48] to remove primer sequences and merge reads using the default parameters. The DADA2 [43] step in the QIIME2 pipeline; merged 16S reads were trimmed to 400bp, followed by dereplication, ASV inference, and chimera removal. The output OTU table was then used to assign taxonomy using a SILVA 138 SSU database [49, 50]. The QIIME2 pipeline outputs a fasta file containing all ASVs with their associated sequences, an OTU table, and a taxonomy table which are used for downstream analyses. Lastly, we removed suspected contaminant ASVs (i.e. those present in sequencing blanks) from both ITS and 16S datasets via the decontam v.1.8.0 package in R v.4.0.3 [44, 51].

### Statistics and other analyses

All downstream analyses were performed in R v4.0.3, unless otherwise noted [44]. To make our data Euclidean, we transformed it with the Hellinger Transformation (*decostand* function) prior to running Principal Component Analysis (PCA, via the *rda* function) [52, 53]. PCA results were plotted using the phyloseq v1.36 function, *plot_ordination* [54]. We used the vegan package v.2.5-7 to calculate diversity metrics and PERMANOVA [52]. For additional statistical analyses and visualization, we used the following in R v4.0.3; TITAN2 v.2.4.1 [55, 56], Venn diagram via eulerr v.6.1.0 [58], network modularity test based on the code from Whitman, *et al* using igraph v.1.2.6 [26, 59], plot visualizations via ggplot2 v.3.3.5 [60], and general data-wrangling via tidyverse v.1.3.0 [61]. The correlation network was calculated using the C++ program FastSpar v.1.0.0 [62], and visualized with the graphic program CytoScape v.3.9.1 [63]. We used Burns, *et al*’s implementation of Sloan’s Community Neutral Model [16, 57] on concatenated 16S and ITS data for each plot. To investigate how neutral model fit changed over time, we divided our samples as equally as possible along the timeseries, and then rarefied samples to 6651 ASVs prior to fitting the neutral model. The fit of the neutral model did not differ substantially over time (Figure S3).

### Data availability

Raw sequencing reads have been submitted to the SRA under accession XXXX. Full results of all statistics and other analyses can be found in the supplemental materials. All code that was used to process and analyze the data is publicly available here: https://github.com/TraxlerLab/BlodgettProject

## RESULTS

### Experimental prescribed fires

To investigate the effect of prescribed fire on soil microbial communities, we established four ten-meter transects within a mixed conifer forest at the University of California’s (UC) Blodgett Forest Research Station, which is around 1360m elevation in the Sierra Nevada Mountains, CA, USA (Figure 1A). Two transects were burned (“Hi” and “Lo”), and the remaining two transects (“1c” and “2c”) functioned as experimental controls that remained unburned throughout a 17 month sampling time series (Figure 1B). The two control plots haven’t burned since the UC acquired the property in 1933. However, prior to that, indigenous people regularly burned the area roughly every 5 - 10 years, up until the gold rush of 1849 and subsequent rapid European colonization displaced and decimated indigenous populations across the region [64]. The Lo plot was treated with low-intensity prescribed fire in 2013 and then again as part of this study in October of 2018. The Hi plot experienced low-intensity prescribed burns in 2002, 2009, and 2017. In January 2019, as part of this study, a slash pile (roughly 20m × 10m × 1.5m) was burned to simulate a higher-intensity prescribed fire (Figure 1B). During the Lo prescribed burn, thermocouples placed 1cm below the soil surface recorded a maximum temperature of 56.0 °C and returned to ambient temperature less than 12 hours after the fire began (Figure 1C). Temperatures 2-6cm below the soil surface did not deviate substantially from the ambient temperature during the Lo burn. During the Hi prescribed burn, thermocouples placed at 3cm below the soil surface recorded a maximum temperature of 71.7 °C, and 12cm below the surface reached a maximum temperature of 52.2 °C. Soil temperature after the Hi burn gradually returned to ambient temperature after four days (Figure 1D). In summary, the heat from the Lo prescribed burn was relatively low, shallow, and not sustained. In contrast, the heat from the Hi prescribed burn was relatively high, deep, and sustained.

**Figure 1:**
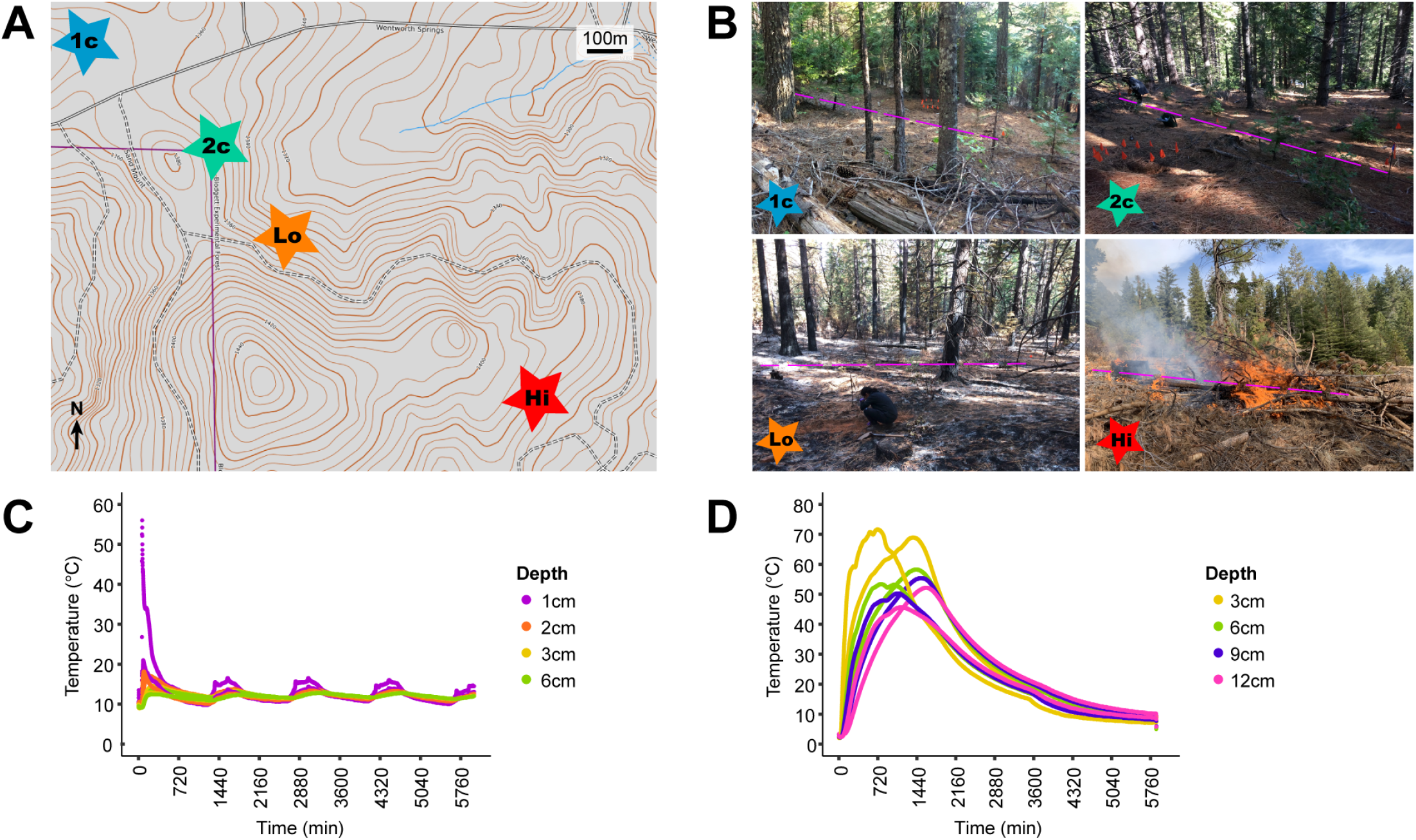
Site description and soil temperature during experimental burns. **(A)** Topographic map showing the location of each transect at Blodgett Experimental Forest. Brown topographic lines denote elevation in meters, solid black lines indicate paved roads, dashed black lines indicate dirt roads, purple line indicates the Blodgett Forest property boundary, and blue lines are streams. “1c” = no burn control #1, “2c” = no burn control #2, “Lo” = low-intensity burn, “Hi” = high-intensity burn. **(B)** Photos each transect. Purple dashed line highlights the location of each 10m transect. Photos of 1c and Lo were taken one week after fire in October 2018. The 2c and Hi transect photos we taken the same day as the start of the fire in January 2019. **(C & D)** Soil temperature measured every 15 minutes starting a few hours prior to each experimental burn near the Lo transect (C), and Hi transect (D). Two thermocouples were placed at each depth, and temperature measurements continued for at least four days, the x-axis is divided into 12-hour increments.

### Prescribed fire alters soil microbial community composition

To investigate how soil microbial communities responded over time to prescribed fire, we collected triplicate soil samples from all four plots prior to burning, and then we continued collecting triplicate soil samples at least once/month, every month (weather permitting) from October 2018 to February 2020. We amplified and sequenced ITS2 and the V3/V4 region of 16S to analyze the soil fungal and bacterial communities, respectively. Principal Component Analysis (PCA) and permutational multivariate analysis of variance (PERMANOVA) found a significant difference between burned and control samples for both bacterial and fungal communities (p < 0.001, Figure 2A). A scatterplot of fungal Principal Component (PC) 1 and PC2 shows three distinct sample clusters associated with either Hi prescribed burn, Lo prescribed burn, or the non-burn controls (Figure 2A). In contrast, a scatterplot of bacterial PC1 and PC2 shows control samples nested within an overlapping region of distinct Hi burn and Lo burn clusters (Figure 2B). For fungal communities, PC1 correlates with soil pH, whereas PC2 correlates with prescribed fire treatment (Figure S4). For bacterial communities, PC1 also correlates with soil pH, however none of our measured environmental or experimental variables clearly explain the variation across PC2 (Figure S5). In contrast to the Hi prescribed burn, the Lo prescribed burn did not affect pH, which highlights that pH is not necessarily the sole determinant of post-fire microbial community assembly. Season, time since burn, average daily air temperature, and precipitation did not explain the variation between our samples (Figure S4 & S5). Prescribed fire generally did not influence diversity or richness over time, with one exception (Figure 2C & S6). After the Hi plot was burned, fungal Shannon diversity, richness, and to a lesser extent, evenness were all reduced post-fire, and remained lower than the Lo, 1c, and 2c plots throughout the sampling time series. Taken together, these results demonstrate that prescribed fire generally had a significant effect on soil microbial community composition, those effects persisted for 17 months post-fire, and higher intensity prescribed fire had a stronger effect on fungi than bacteria.

**Figure 2.**
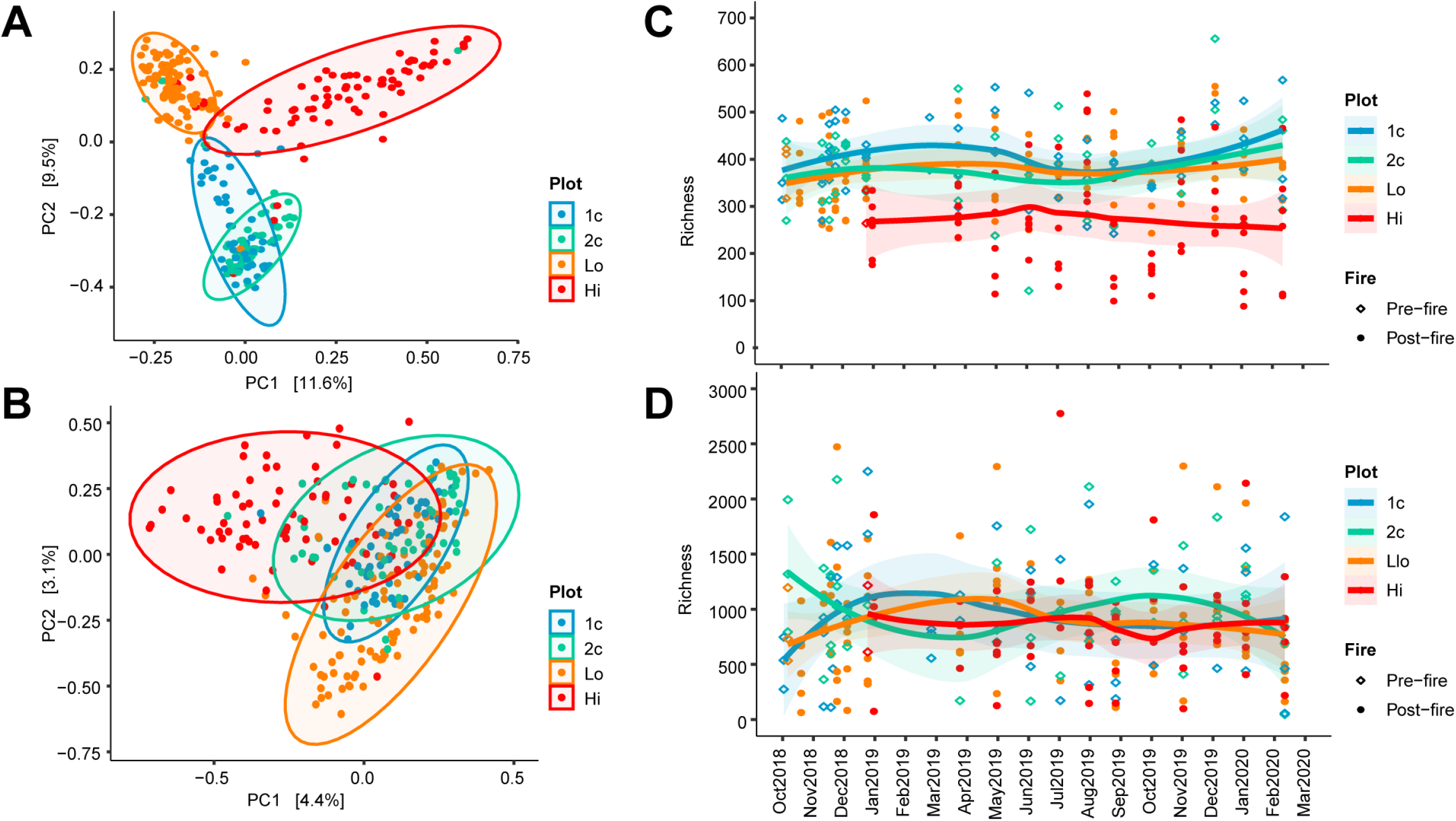
Fire alters soil microbial community structure. **(A & B)** Principal Component Analysis (PCA) on Hellinger-transformed ITS (A) & 16S (B) amplicon community sequencing data. Axes are the two Principal Components (PC) that explained the most variation in the data (% variation noted on each axis). Ellipses = 95% confidence interval. PERMANOVA p < 0.001 for burned vs. unburned in both ITS and 16S. **(C & D)** Community richness over time for ITS (C) and 16S (D) data. Points are individual samples, and the data for each plot are summarized by fitting a local polynomial regression line. The shaded area around each line indicates a 95% confidence interval.

### Indicator taxa associated with changes in community composition following prescribed fire

To identify significant shifts in community composition over time, we conducted a Threshold Indicator Taxa Analysis (TITAN). TITAN combines change point analysis [64, 65] and indicator species analysis [66] to find time-points where community composition changed significantly, while also highlighting dynamic taxa that were indicators of whole-community shifts [55, 56]. TITAN identified two significant change points along the time series for each of our four plots (Figure 3A and S7). The first change point in each plot is associated with indicator taxa that generally decline in abundance over time (“negative responders”), and the second change point is where indicators tend to increase over time (“positive responders”) (Figure 3). The negative change points for both burned plots occurred immediately following prescribed fire, and then roughly one year later. In contrast, change points in our control plots occurred during winter months (Figure 3A). For example, plot 1c possibly experienced community shifts associated with minimum mid-winter temperatures, while 2c possibly experienced community shifts associated with the first precipitation event of the wet season (Figure S8). TITAN identified substantially more indicator taxa in burned plots (Lo = 156, Hi = 121) than in control plots (1c = 33, 2c = 73). A majority of control plot indicators were negative responders (1c: 88%, 2c: 64%), whereas burned plot indicators were roughly evenly distributed between negative and positive responders (Figure 3B).

**Figure 3.**
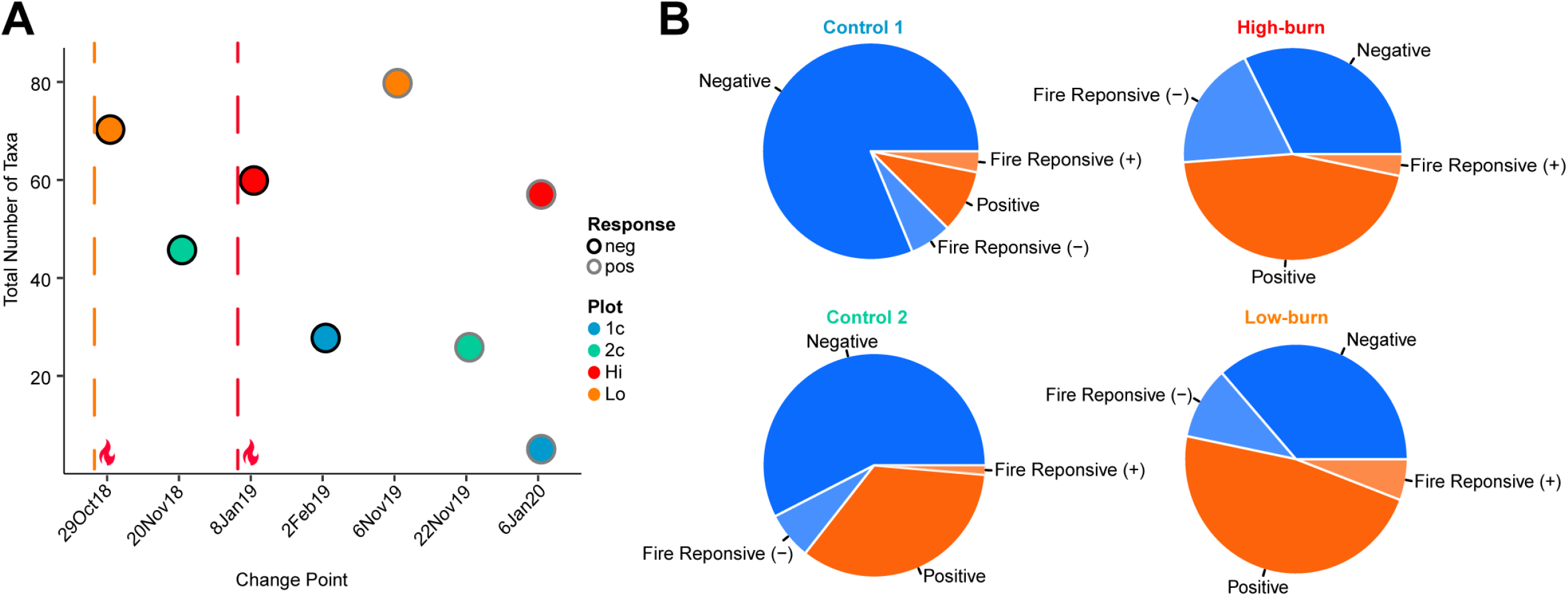
Shifts in microbial community composition associated with fire and TITAN identified a greater number known fire-responsive taxa as indicators in fire-treated plots. **(A)** Total number of indicator taxa identified by TITAN as either positively or negatively responsive, and the corresponding change point date for each plot. Dashed vertical line and flame emoji denote the time-point at which burned plots were burned. **(B)** Proportion of known fire-responsive taxa within positive (red) or negative (blue) response groups for Control 1, Control 2, Hi-burn, and Lo-burn plots.

TITAN indicator taxa in the burned plots include many genera that have been previously described as fire-responsive (Figure 3B). We compiled a list of microbial genera from previous literature that responded positively to fire (File S2). We highlighted taxa assigned to these genera as lighter-hued sections in Figure 3B. Notably, this included the fungal genera *Pyronema, Geopyxis, Anthracobia, Lyophyllum*, and *Myxomphalia*, and the bacterial genera *Massilia, Flavobacterium*, and *Blastococcus*. Taken together, these findings demonstrate that fire led to both immediate and delayed effects on soil microbial community composition, and dynamic responders to fire included previously documented fire-associated genera.

### Prescribed fire intensity drives microbial community substructure

To identify if there was significant network substructure in our community, we first calculated a correlation network (via FastSpar, which is based on Pearson correlation), and then used Clauset, *et al*’s fast greedy clustering method to test for network substructure [67]. Clauset, *et al*’s test for community substructure, or modularity, resulted in a Q-value of 0.329, and Q > 0.3 is considered a good indicator of network modularity [67, 68]. The network was composed of a total of 19 modules, but 98.6% of all taxa fell within the first three modules (Supplemental File S3), thus we focus on Modules 1-3. To understand factors that might underlie this modularity, we compared module assignment to relative abundances in each plot and to TITAN results (Figure 4). We set a conservative threshold to determine if a taxon was highly abundant in each sample; greater than the 95^th^ percentile for all abundance values. 41% of all taxa in the network were consistently below the 95^th^ percentile, thus not highly abundant (black in Figure 4A). 24% of all taxa were highly abundant in both prescribed burn and control plots, thus uninformative (gray in Figure 4A). 22% of all taxa were highly abundant in burned plots, but not control plots, and 70% of these burn-abundant taxa were assigned to network Modules 2 and 3 (red in Figure 4A). In contrast, the remaining 13% of all taxa were highly abundant in control plots, but not burned plots (blue in Figure 4A), and 77% of these control-abundant taxa were assigned to Module 1 (Figure 4A). A similar pattern was maintained when the network was filtered for TITAN indicator taxa; Module 1 was enriched for indicators of the control plots (44% controls, 38% Lo, 16% Hi) and Modules 2 & 3 were dominated by prescribed burn indicators. Notably, indicators of the Hi prescribed burn largely fell within Module 2 (65% Hi, 17% Lo, 18% controls), whereas Lo prescribed burn indicators were predominantly assigned to Module 3 (71% Lo, 7% Hi, 22% controls). Taken together, these results point toward cohorts of microbial taxa that responded positively (Modules 2 & 3), or negatively to fire (Module 1). This network substructure further differentiated between taxa that were indicators for either the low-intensity burn treatment or high-intensity burn treatment, indicating that the soil microbial community responded uniquely to fires of differing intensities.

**Figure 4.**
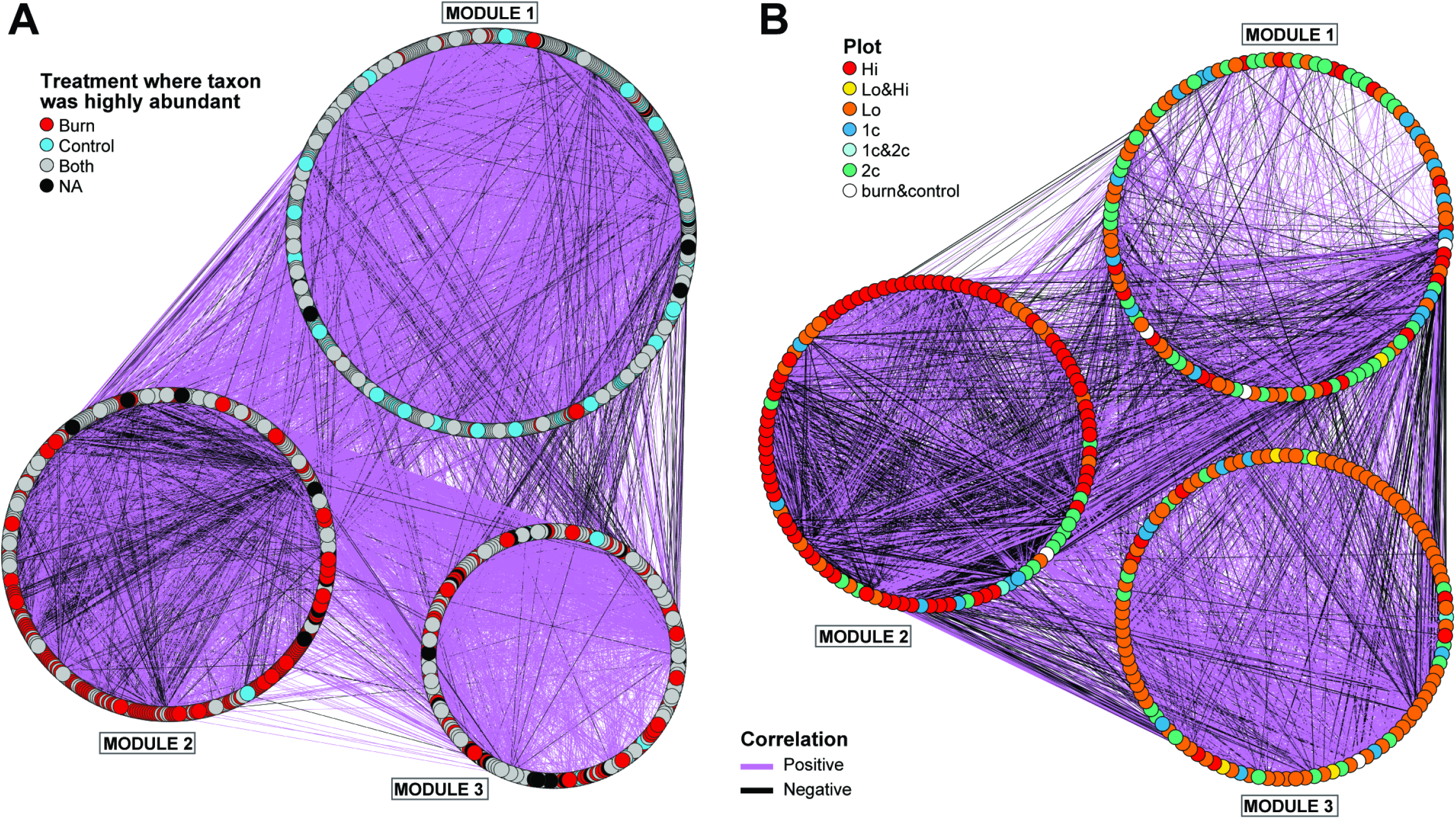
Correlation network clusters taxa by fire treatment. Correlation network arranged into modules defined by greedy clustering [67], which identified a total of 19 modules, modules with less than 10 nodes were excluded. Nodes are taxa, and lines represent a significant correlation between taxa (p < 0.01, FastSpar). Purple line = positive correlation, black line = negative correlation. **(A)** Complete network. For clarity, only edges and nodes with correlations > 0.3 are shown. Node colors indicate the treatment(s) in which the taxon was highly abundant; red = burn, cyan = control, grey = both burn and control, and black indicates taxa that were consistently in low abundance (< 95th percentile) and subsequently excluded from the presence/absence analysis. **(B)** Subset of the complete network showing the 335 taxa that were identified as significant indicators by TITAN. All significant correlation values are shown (p < 0.01), and line width is proportional to the correlation value. Node colors indicate in which plot the taxon was identified by TITAN as an indicator, briefly, cool colors represent control plots and warm colors represent burned plots.

### A combination of neutral and deterministic processes drive soil community assembly patterns

To illuminate broad ecological processes driving community assembly, we investigated the contribution of neutral and deterministic processes in structuring the soil microbial community. We fit the Sloan Neutral Community Model to our data [16, 57], and then examined the taxa that fell within (neutral) or outside (deterministic) the neutral prediction (Figure 5). On average 92.8% (± 0.7%) of taxa in each of our plots fit the neutral prediction (Figure 5A), and the neutral model fit was roughly equivalent across all plots (R^2^ = 0.63 - 0.73) (Figure 5B). Importantly, the subsets of taxa identified by both TITAN and the correlation network analysis were enriched for taxa that fell outside the neutral prediction (Figure 5A). Roughly half of all taxa identified as indicators by TITAN were non-neutral, and 54% of taxa in network modules 1 - 3 were also nonneutral. Notably, Module 2 was enriched for taxa that were non-neutral in the Hi burn plot, whereas Module 3 was enriched for non-neutral taxa in the Lo burn plot. Several previously described fire-responsive taxa fell outside the neutral prediction in both burned plots, which are highlighted in Figure 5B. Lastly, Sloan’s Neutral Community Model estimates the rate of dispersal, or migration (m) into the community. This migration value in all plots was close to zero (m < 0.01), indicating that dispersal likely has very limited influence on community structure in either control or burned plots. Together these data demonstrate that a majority of taxa in our samples are present due to stochastic, or neutral processes. However, the neutral model did not predict the presence of the majority of indicator taxa that dynamically responded to fire, implying that their presence was likely the result of deterministic processes, i.e. selection.

**Figure 5.**
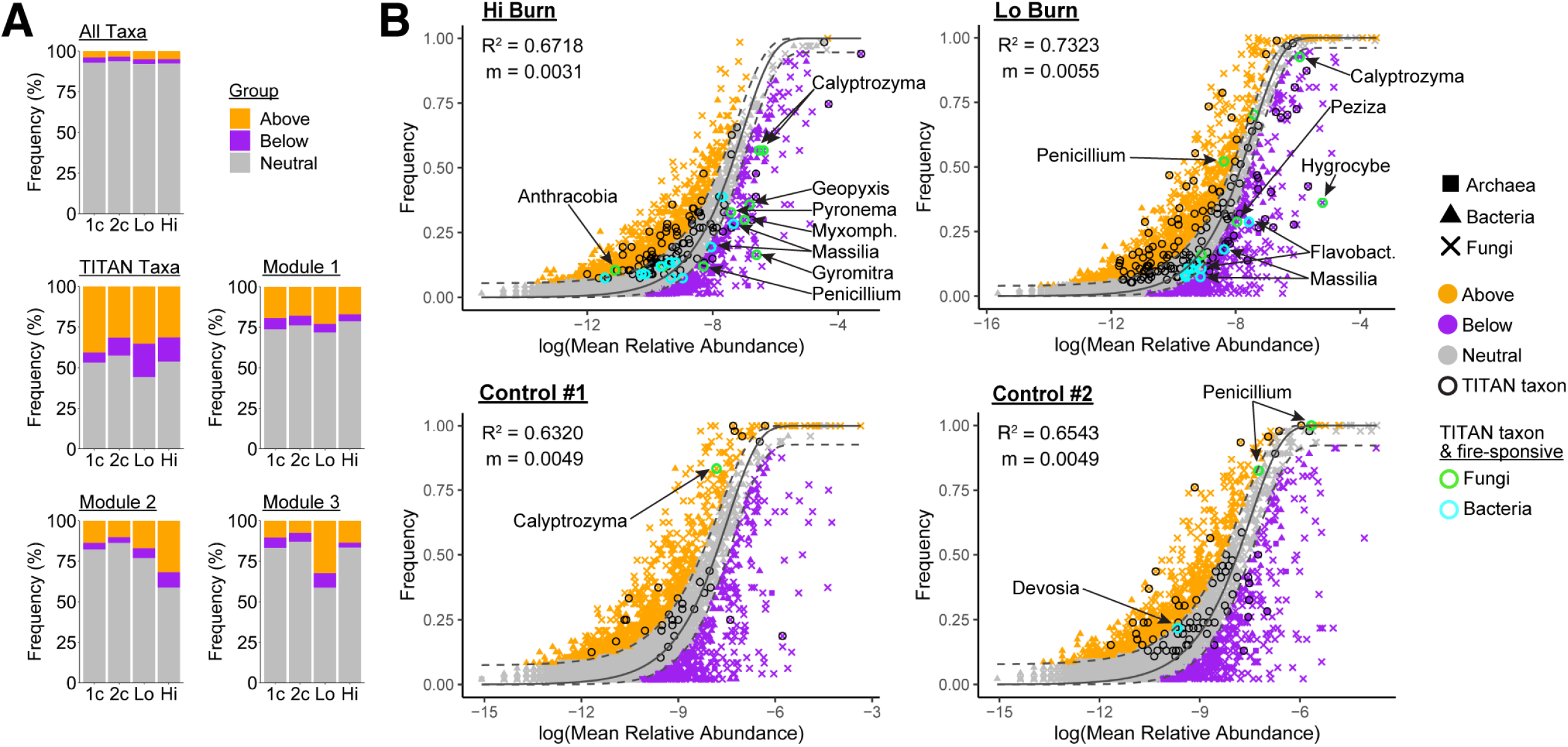
A combination of neutral and deterministic processes drive community assembly patterns. **(A)** Stacked bar-plots illustrating the number of taxa that fell within, above, or below the neutral model in each plot, for all taxa and subsets of taxa based on TITAN or the correlation network analysis. **(B)** The Sloan Neutral Community Model fit to ITS and 16S data in each plot. The neutral prediction is a solid dark-grey line, with 95% confidence interval around indicated with dashed dark-grey lines. Taxa that fall within this 95% confidence interval are colored grey. Taxa that fall outside the neutral predicted are colored either orange (above) or purple (below). Taxa that were found to be indicators via TITAN are circled in black, and of these indicators, those that have been previously described as fire-responsive are circled in either green (fungi) or cyan (bacteria). Fire-responsive TITAN indicators that fell outside the neurtral prediction are named. R^2^ quantifies how well the neutral model fit the data, and m is the estimated migration rate.

### Pyrophilous genera associated with prescribed fire

We leveraged the results of TITAN, neutral model fit, and the correlation network analyses to independently identify pyrophilous taxa within our data (Figure 6). Figure 6A-C details our methodology for identifying pyrophilous taxa. Briefly, we subset the correlation network for the 96 taxa that were unique to burned plots as TITAN indicators and/or non-neutral (Figure 6A&B), and then further filtered this list for taxa that were highly abundant only in burned plots (Figure 6C). These filtering steps resulted in nine unique genera, which we consider to be pyrophilous: the fungi *Pyronema, Geopyxis, Lyophyllum, Myxomphalia, Rhodosporidiobolus*, and the bacteria *Massilia, Bacillus, Flavobacterium*, and *Cellvibrio*. These taxa are highlighted in the network in Figure 6B, and we illustrate their average normalized relative abundance over time in Figure 6D. All pyrophilous genera except *Cellvibrio* were assigned to network Module 2. *Cellvibrio* fell within Module 3, along with one of three *Flavobacterium* ASVs (Figure 6B). The average abundance of the pyrophilous genera over time in both the Lo burn and control plots generally mirrors precipitation events (Figure S8), however abundances were substantially higher after burning and decreasing over time. *Pyronema* dominated immediately after the Lo prescribed burn prior to any precipitation (the Lo plot was burned on 16 Oct 2018 and the first rain was on 20 Nov 2018). Following the start of the rainy season, *Pyronema* and all the pyrophilous bacteria peaked in abundance (Figure 6D). The Hi plot was treated with prescribed fire during a mid-winter dry period, and precipitation occurred one day after starting the fire. *Geopyxis* peaked in abundance immediately after the Hi burn (Figure 6D). *Lyophyllum, Bacillus*, and *Massilia* also experienced a peak in abundance at the first sampling time point following the Hi prescribed burn. In contrast to the Lo burn, *Pyronema* peaked in abundance three months after the Hi burn, along with *Rhodosporidiobolus, Flavobacterium*, and *Massilia*. All nine pyrophiles increased dramatically in abundance following the Hi prescribed burn. In contrast, in the Lo plot, *Lyophyllum* and *Rhodosporidiobolus* were not abundant at any time point. In conclusion, this combined analysis independently identified nine pyrophilous genera that responded strongly and positively, likely through deterministic processes associated with prescribed fire at Blodgett Forest.

**Figure 6.**
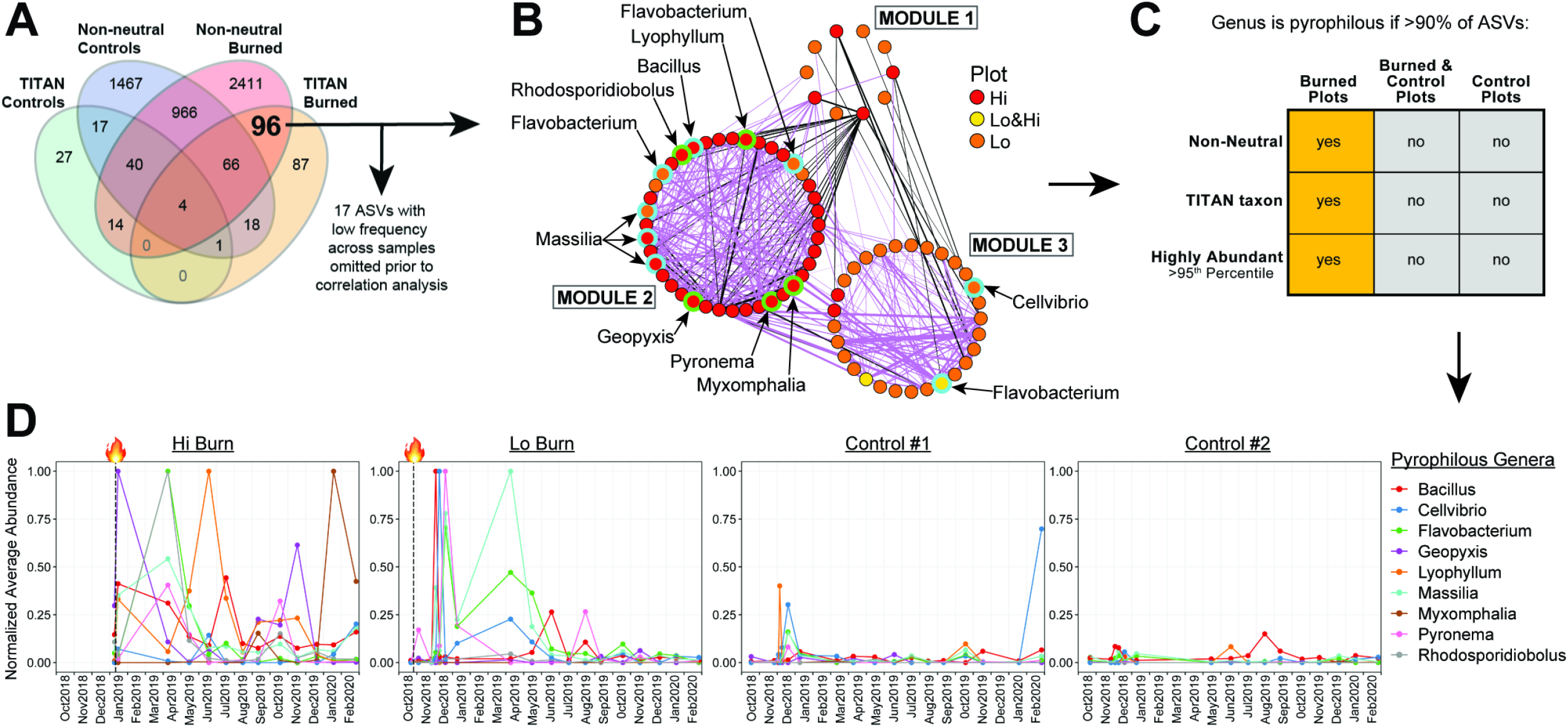
Identification of pyrophilous genera at Blodgett Forest, and their dynamics over time. **(A)** Venn Diagram showing the number of taxa that were either identified as indicators by TITAN or fell outside the neutral prediction. **(B)** Filtered version of the correlation network depicted in Figure 4. This network was filtered for the 96 ASVs that were unique to burns in A. Pyrophilous taxa are highlighted and labeled (green outline = fungus, blue outline = bacteria). Lines thickness it proportional to the correlation value (purple = positive, black = negative). Node colors indicate the plot in which the taxon was an indicator. **(C)** Diagram describing the criteria for defining pyrophilous genera in our dataset. **(D)** Relative abundance of pyrophilous genera over time in each plot. Dashed vertical line and flame emoji denote the time-point at which burned plots were burned. Each point represents the average abundance of all ASVs for each genus at each timepoint, and average abundances were normalized to the maximum average abundance for each genus.

## DISCUSSION

As the incidence of wildfire increases with ongoing climate change, strategies for mitigating their impact, including prescribed burns, are becoming increasingly relevant. Here, we sought to understand the impact of prescribed fire on soil fungal and bacterial communities, and the ecological processes that influence post-fire microbial community assembly. We undertook extensive time-course sampling of two prescribed burn plots, one which experienced a low intensity fire, and a second that experienced a higher intensity fire. We found that fire significantly altered both the fungal and bacterial communities in the burned plots, as compared to unburned control plots. Using a series of downstream analyses, we conclude that, while a large portion of the post-prescribed burn community is likely assembled via neutral processes, a subcommunity that includes pyrophilous organisms likely arises through deterministic processes. A number of other studies have begun to paint a picture of post-fire microbial community structure [12, 22, 24, 26, 31, 69–73], and this work further builds on those by identifying members of the subcommunity that deterministically respond to fire. In doing so, this work lays the foundation for building a process-driven understanding of microbial community assembly in the context of the classical disturbance regime of fire.

Neutral processes such as passive dispersal and ecological drift have been shown to be important during the colonization of unoccupied environments, and a recently burned landscape may seem, superficially, like such an environment. However, many organisms survive below the soil surface even during intense wildfires [12], and organismal survival becomes more likely with decreasing fire intensity. This notion is consistent with our data showing that lower-intensity prescribed fires had a minimal impact on richness (Figure 2C-D). We also found that the majority (∼93%) of the soil microbial community at Blodgett Forest could be accounted for by a model of neutral assembly (regardless of fire occurrence). Thus, the community structure in our burned plots likely reflects a combination of; (1) the legacy of neutrality in the soil microbial communities that was present prior to fire, (2) *de novo* community assembly through neutral processes, and (3) selection of a subcommunity of microbes adapted to postfire environments.

To identify members of the fire-responsive subcommunity, we used TITAN and a correlation network analysis, in conjunction with a neutral model. TITAN highlighted individual indicator taxa whose abundances were dynamic across time, while correlation network analysis identified sub-groups of taxa whose members were linked through correlations in abundance, irrespective of time. Importantly, the subcommunity identified by the combined correlation analysis and TITAN was enriched for non-neutral taxa (Figure 5A), indicating that deterministic processes likely played an important role in the assembly of this fire-responsive subcommunity. Fire is a dramatic selective force that has a myriad of effects on soil, such as direct heating during the fire and enduring post-fire effects including increased pH, increased hydrophobicity, decreased bioavailability of nutrients (especially nitrogen), and the deposition of a layer of PyOM and mineral ash. Our data do not allow us to directly distinguish between these possible selective forces associated with prescribed fire. However, it is notable that in our Lo burned plot we did not observe any significant changes in pH, and there was little-to-no effect of heat below the soil surface (Figure 1C, S4 & S5), yet we still observed a significant effect on community composition. Thus, we hypothesize that factors beyond temperature and pH, such as deposition of PyOM, may underlie assembly of the fire-responsive sub-community following low-intensity fire. Further investigation will be required to test this hypothesis *in situ*.

Taxa that fall above the neutral prediction in Sloan’s Neutral Community Model are found more frequently than would be predicted by their abundance in the metacommunity. Conversely, taxa that fall below are highly abundant in fewer samples than would be expected, resulting from a patchy distribution across samples. Burns, *et al* suggested that taxa above the neutral prediction are being selected for, whereas taxa below may be seen as ‘invasive’, or are dispersal limited, or are selected against. It’s notable that most taxa that we identify as pyrophilous fell below the neutral partition in our burned plots (Figure 5B), indicating that they were found at high abundance with relatively low frequency. Such a patchy distribution may be explained by the fact that the effects of fire across a landscape result in a spatially heterogeneous mosaic [74, 75]. For example, individual soil patches experience varied levels of perturbation from irregular heating, as well as non-uniform deposition of PyOM and changes in pH. Alternatively, patchy distributions of locally abundant taxa can arise as a result of interference competition that leads to spatial segregation of competing organisms [76, 77], although more study will be required to link this possibility to taxa that fall below neutral model predictions.

While there are numerous previous descriptions of pyrophilous taxa (File S2), we add to this body of work by using multiple analyses to independently identify pyrophilous taxa in post-prescribed burn environments. Specifically, we included taxa that; (1) were dynamic over time in burned plots according to TITAN, (2) showed strong correlation patterns with other community members, (3) fell outside the neutral expectation in burned plots, and (4) were only highly abundant in burned plots (Figure 6A-C). Only nine genera passed through this stringent filtering process, six of which have been previously reported as pyrophilous (*Pyronema, Myxomphalia, Lyophyllum, Geopyxis, Massilia*, and *Flavobacterium*). The three genera that are currently absent from other pyrophile literature are *Bacillus, Cellvibrio*, and *Rhodosporidiobolus. Bacillus* (Firmicutes) are well-known, ubiquitous soil bacteria that form remarkably resistant spores, which could be important for surviving the heat of intense fires. Beyond survival, we note that several species of *Bacillus* are known to degrade polycyclic aromatic hydrocarbons, which may be relevant in the consumption of PyOM [78, 79]. *Cellvibrio* (Gammaproteobacteria) are also common soil inhabitants that have recently drawn biotechnology interest for their production of xylanases and other carbohydrate active enzymes [80], and some isolates have demonstrated the ability to fix nitrogen [81], which could be critically important for life in nitrogen-depleted post-fire soils [82]. *Rhodosporidiobolus* is a red yeast in the Pucciniomycotina (Basidiomycota) that was first described in 2015 [83] and has since been found to increase in abundance after biochar addition to tea orchard soil [84], and demonstrated a robust ability to degrade lignin and other aromatic compounds [85]. These examples point toward the notion that *Rhodosporidiobolus* may be able to utilize the PyOM component of a recently burned environment, similar to *Pyronema domesticum* [86].

All microbial communities are assembled through some combination of the deterministic process of ecological selection (encompassing environmental filtering and biotic interactions) and the stochastic processes of passive dispersal, ecological drift, and mutation/speciation. Here we used parallel analyses to identify taxa that were temporally dynamic, showed patterns of co-occurrence, and whose presence was likely attributable to deterministic processes. In delineating this pyrophilous subcommunity, this work lays the foundation for future investigations into the mechanisms that drive pyrophilous community assembly in post-fire environments. Furthermore, we speculate that interactions among members of this sub-community may impact the degradation and reintegration of pyrolyzed organic matter in areas under pressure from increasingly frequent wildfires. Finally, this work sets the stage for understanding the role of this subcommunity in stimulating the recovery of the broader community of micro-and macro-organisms, while also providing a starting point for future studies aimed at harnessing deterministic interventions to enhance community resilience to fire.

## Supporting information

FischerPatel_SupplementalFigures

## ACKNOWLEDGEMENTS

This research would not have been possible without the incredible support of Ariel Thomson Roughton, Robert York, Amy Mason, and everyone at Blodgett Forest Research Station. We are also grateful for Peter Wyrsch, Thea Whitman, and Luis Cantu Morin for many fruitful conversations. This research used the Savio computational cluster resource provided by the Berkeley Research Computing program at the University of California, Berkeley (supported by the UC Berkeley Chancellor, Vice Chancellor for Research, and Chief Information Officer). The sequencing was carried out at the DNA Technologies and Expression Analysis Cores at the UC Davis Genome Center, supported by NIH Shared Instrumentation Grant 1S10OD010786-01. This research was supported by the DOE Office of Science and Office of Biological and Environmental Research (BER); grant no. DE-SC0020351 to MT.

