## Supplementary material for "Prescribed fire selects for a pyrophilous soil subcommunity in a northern California mixed conifer forest": FischerPatel_SupplementalFigures

**Table of Contents**

Figure S1……………………………………………………………………………………...3
Figure S2……………………………………………………………………………………...4
Figure S3……………………………………………………………………………………...5
Figure S4……………………………………………………………………………………...6
Figure S5……………………………………………………………………………………...7
Figure S6……………………………………………………………………………………...8
Figure S7……………………………………………………………………………………...9
FigureS8…………………………………………………………….……………………11

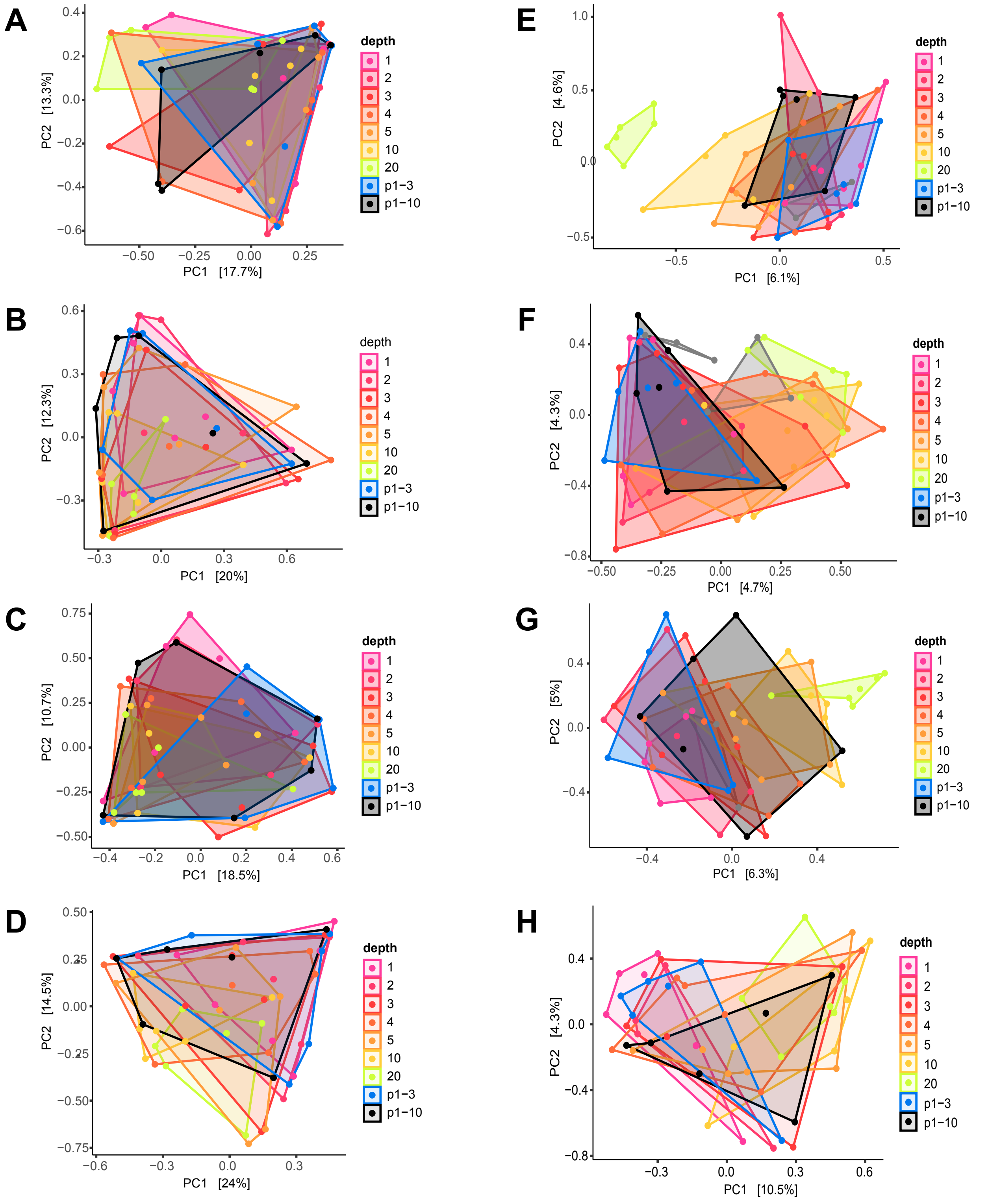


**Figure S1. Effect of depth on soil microbial community structure.** PCA plots examing the effect of depth on ITS (A-D) and 16S (E--H) community structure in a control versus a burned plot, sampled at two timepoints, pre- and post- fire. ITS community in burn plot pre- (A), post- (B) fire and in control plot pre- (C), post- (D) fire. 16S community in burn plots pre- (E), post- (F) and in control plot pre- (G), post- (H) fire. There is no significant difference between depths (PERMANOVA $p > 0.05$, n=6)

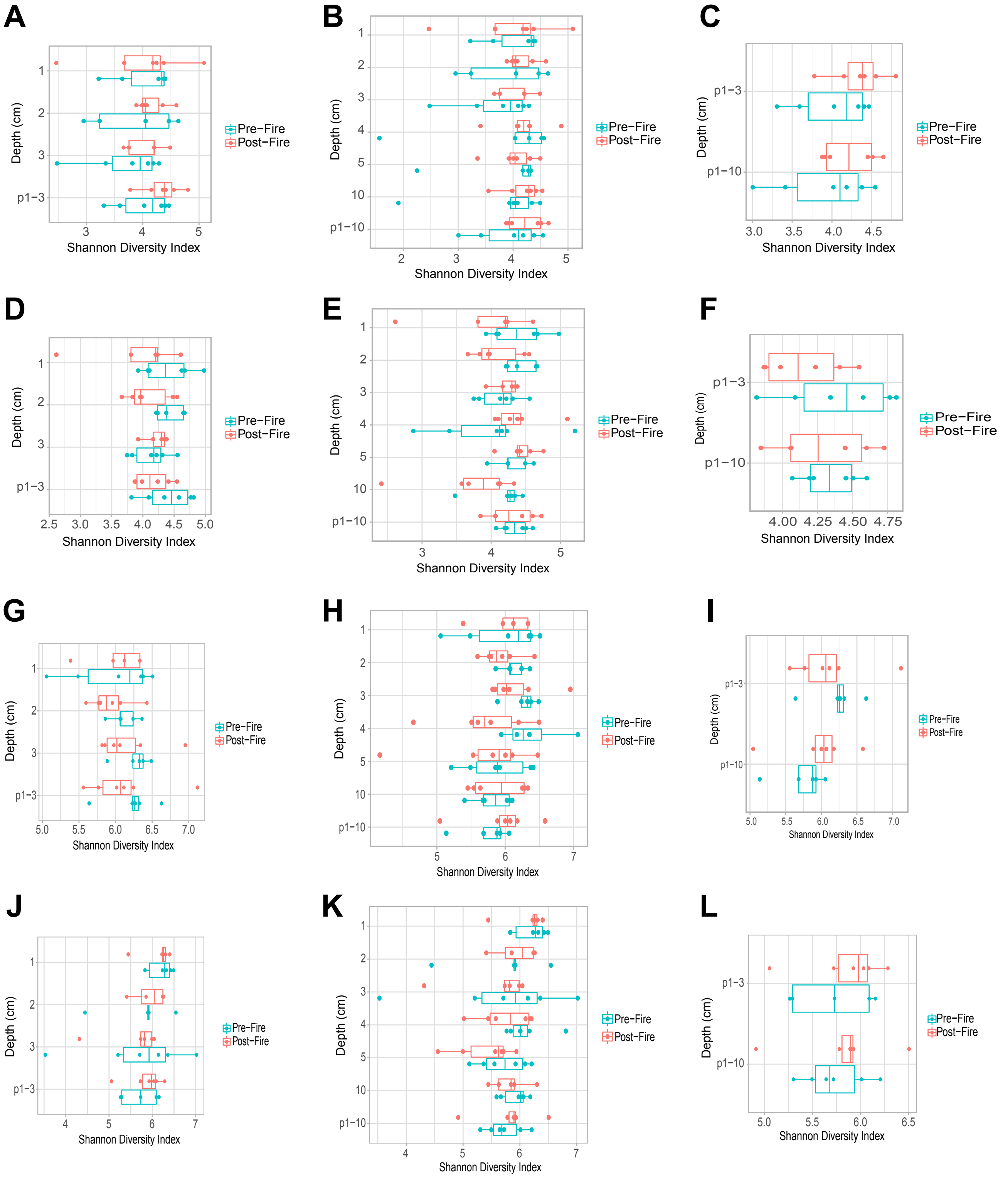


**Figure S2. Shannon Diversity indices by depth in pre- and post- fire treatment.** No significant differences were detected in Shannon Diversity in either ITS (A-F) and 16S (G-L) communities, based on ANOVA and t-test by depth (n=5). ITS community in treatment plots for 1-3cm (A), ANOVA = 0.742 pre-fire, 0.611 post-fire. 1-10cm (B), ANOVA = 0.995 pre-fire, 0.948 post-fire. Pooled 1-3cm and 1-10cm (C), t-test = 0.7613 pre-fire, 0.5872 post-fire. ITS community in control plots for 1-3cm (D), ANOVA = 0.444 pre-fire, 0.673 post-fire. 1-10cm (E), ANOVA = 0.562 pre-fire, 0.104 post-fire. Pooled 1-3cm and 1-10cm (F), t-test = 0.7495 pre-fire, 0.4787 post-fire. 16S community in treatment plots for 1-3cm (G), ANOVA = 0.572 pre-fire, 0.773 post-fire. 1-10cm (H), ANOVA = 0.156 pre-fire, 0.707 post-fire. Pooled 1-3cm and 1-10cm (I), t-test = 0.187 pre-fire, 0.746 post-fire. 16S community in control plots for 1-3cm (J), ANOVA = 0.634 pre-fire, 0.322 post-fire. 1-10cm (K), ANOVA = 0.652 pre-fire, 0.256 post-fire. Pooled 1-3cm and 1-10cm (L), t-test = 0.187 pre-fire, 0.589 post-fire.

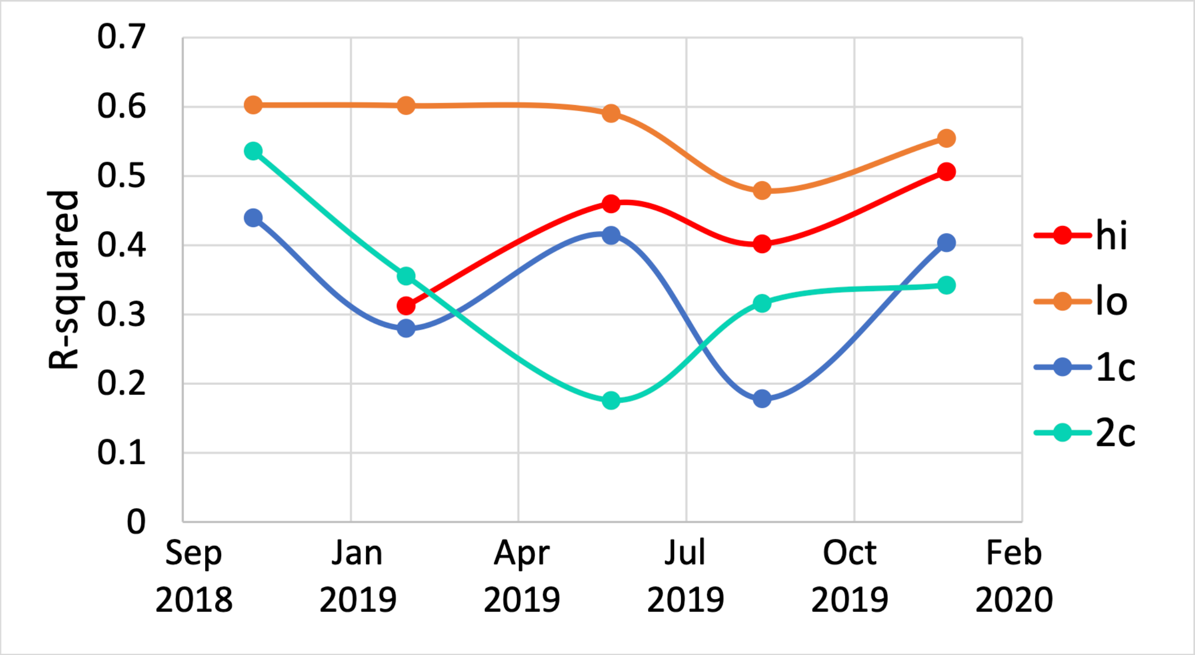


**Figure S3. Neutral Model fit over time.** To compare across plots, samples were divided into roughly even groups along the sampling timeline such that each group contained 3-5 timepoints (x=midpoint) and 9-27 samples, and taxa were rarefied to 6651 ASVs.

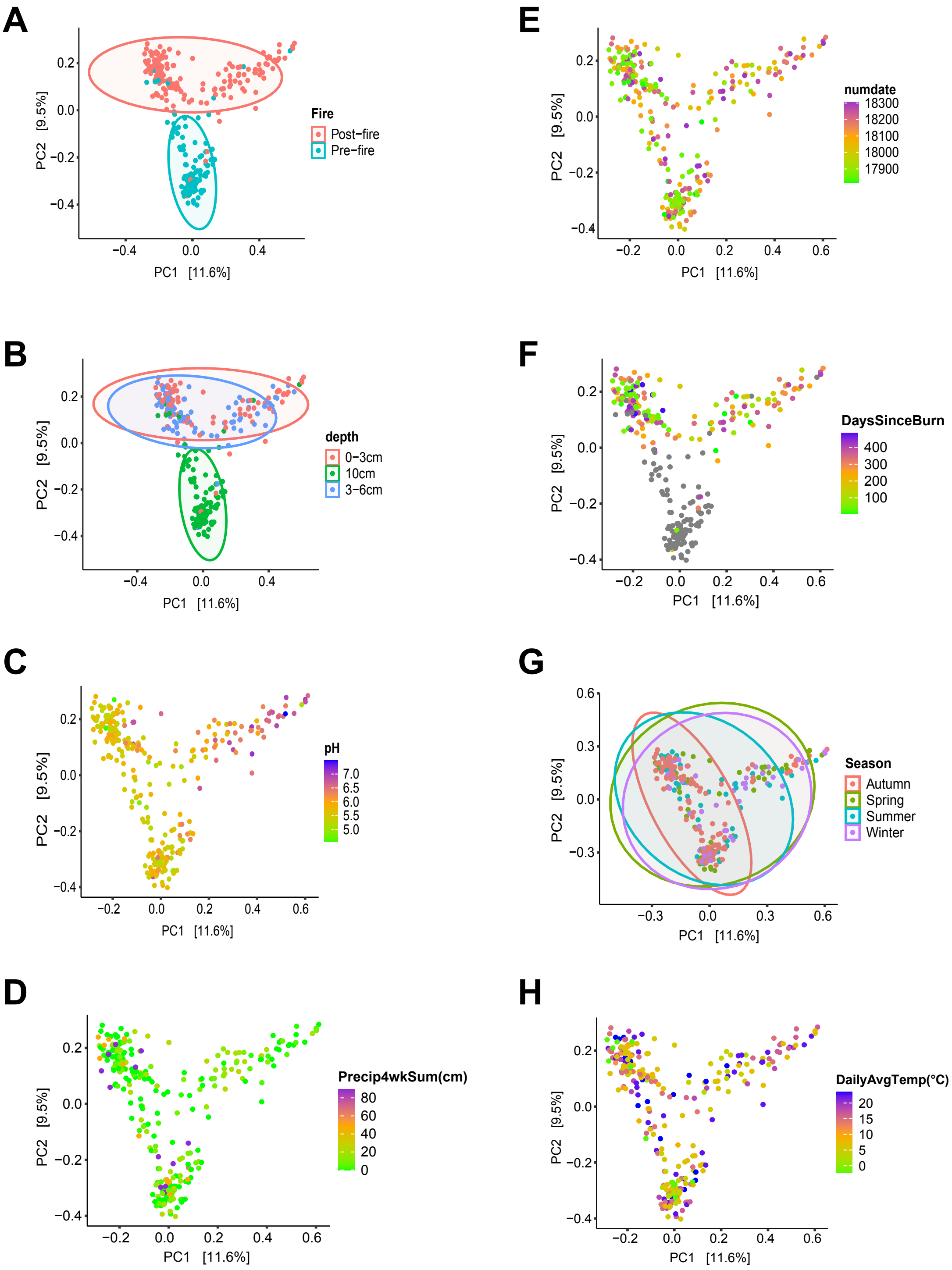


**Figure S4. ITS community PCA plots.** PCA plots examining the effect of fire (A), depth (B), pH (C), precipitation (D), date (E), days post-fire (F), season (G) and average daily temperatures (H) on the ITS community.

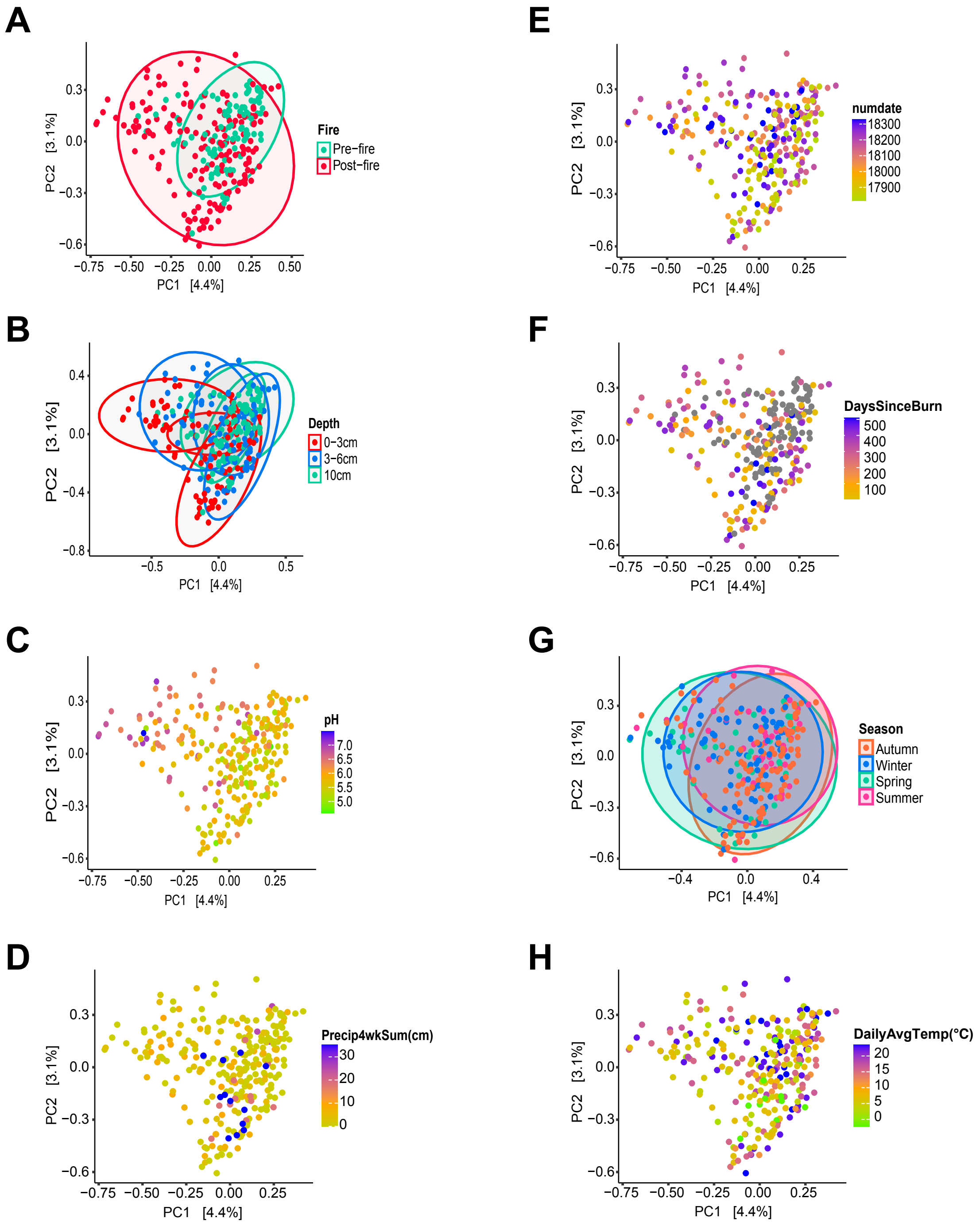


**Figure S5. 16S community PCA plots.** PCA plots examining the effect of fire (A), depth (B), pH (C), precipitation (D), date (E), days post-fire (F), season (G) and average daily temperatures (H) on the 16S community.

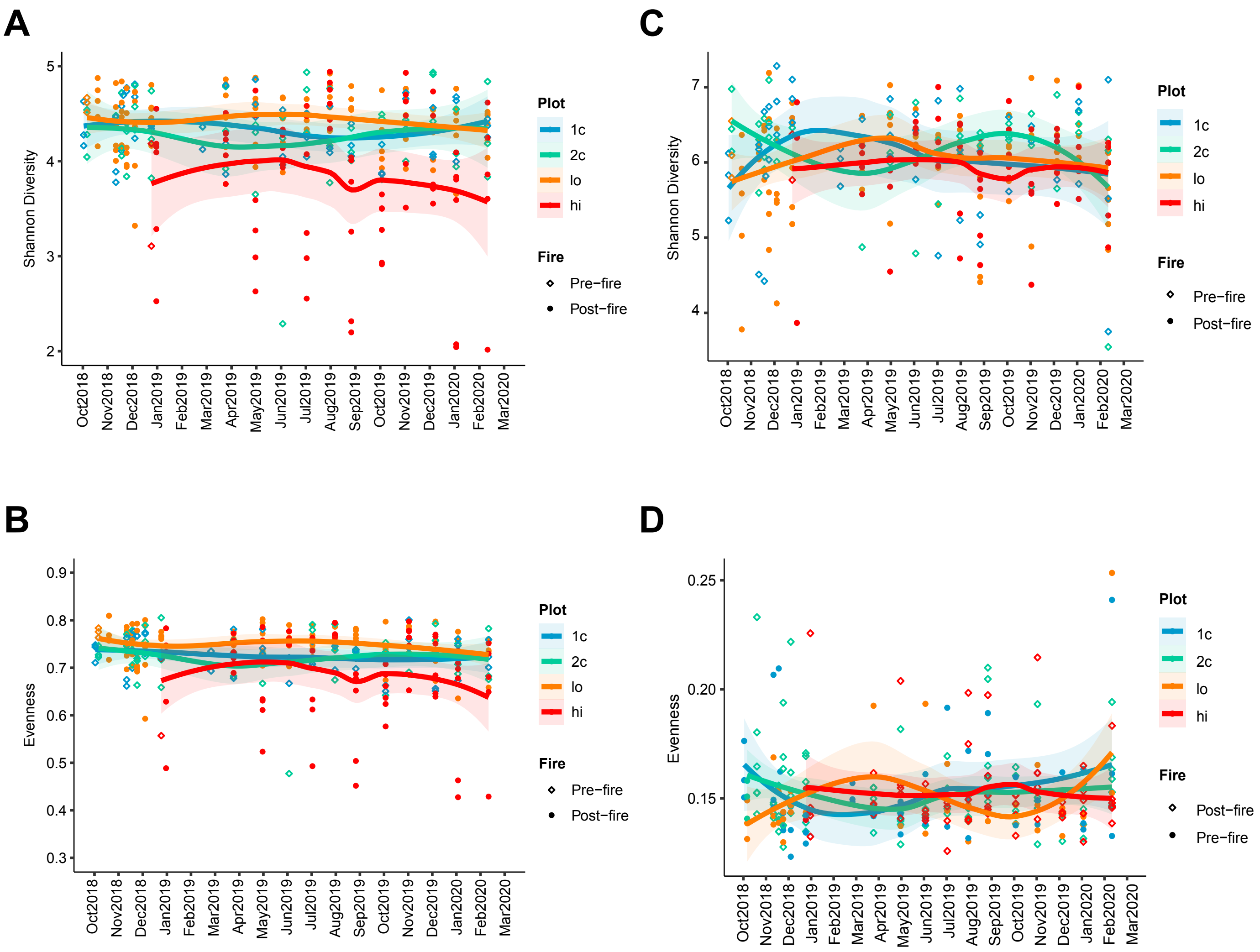


**Figure S6. Shannon Diversity and community evenness.** Shannon Diversity indices over time in both control and treatment plots for the ITS (A) and 16S (C) communities. Community evenness over time in both control and treatment plots for the ITS (B) and 16S (D) communities.

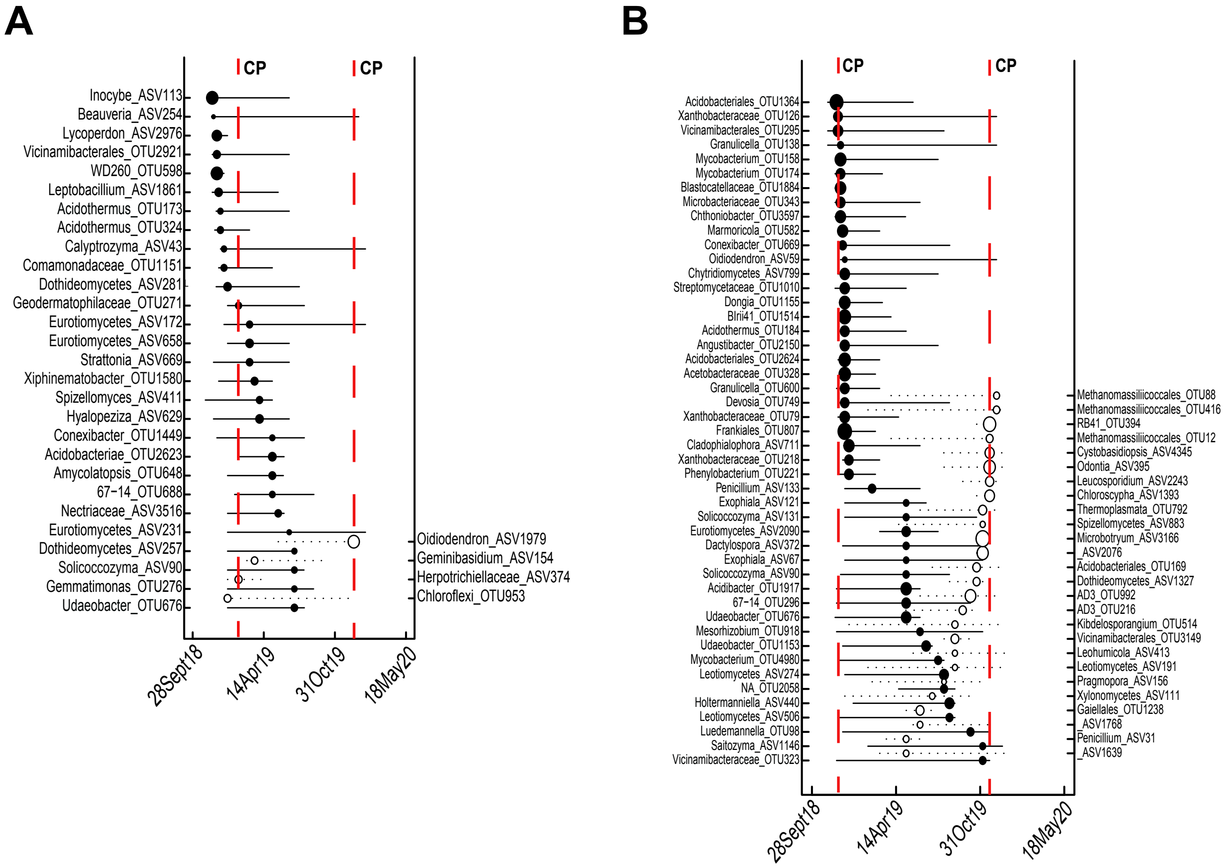


**Figure S7. TITAN.** Full TITAN plots with all identified indicator taxa and thier associated z-scores for control 1 (A) and control 2 (B).

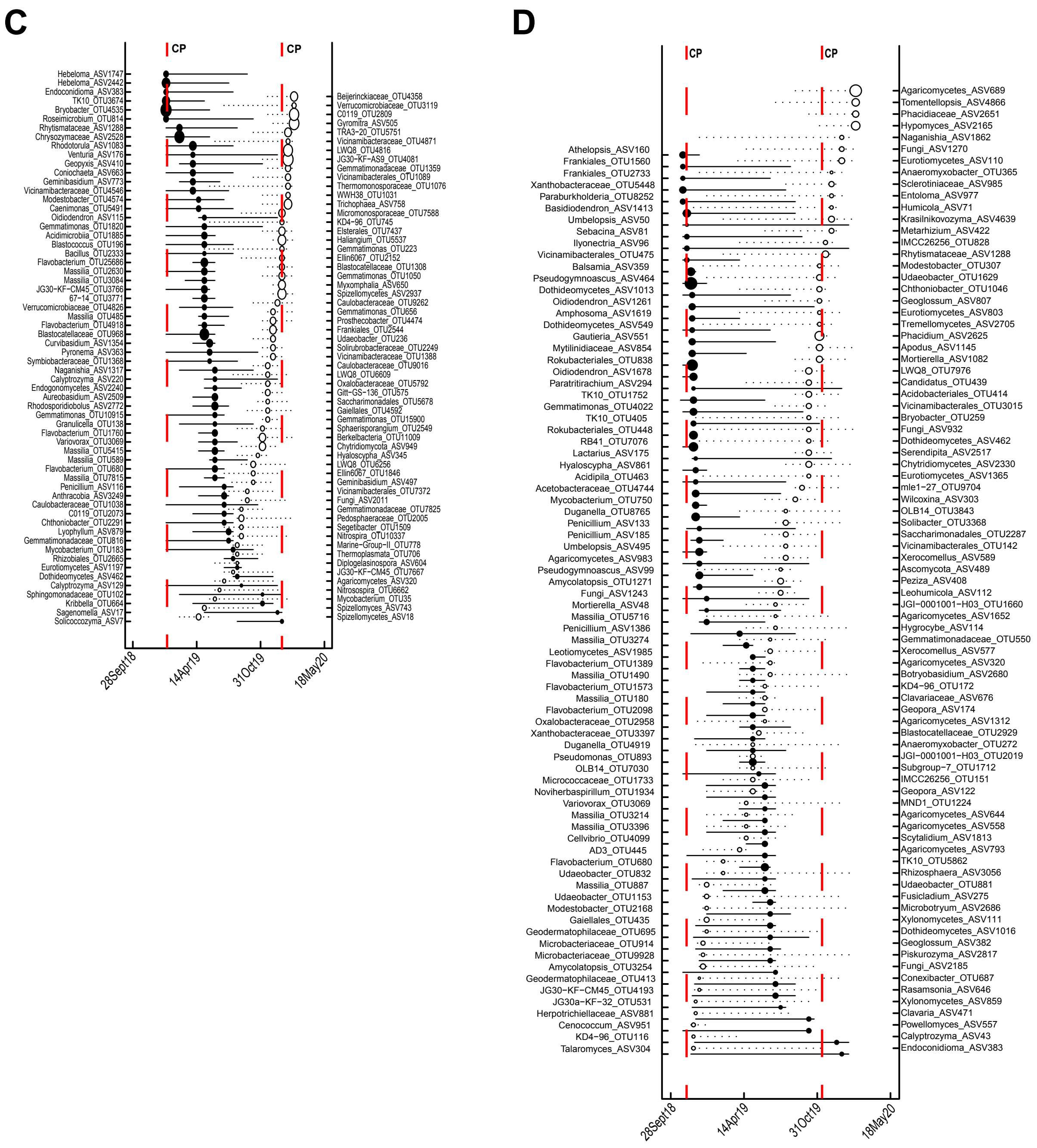


**Figure S7. TITAN.** Full TITAN plots with all identified indicator taxa and thier associated z-scores for low-burn (C) and high-burn (D) treatments.

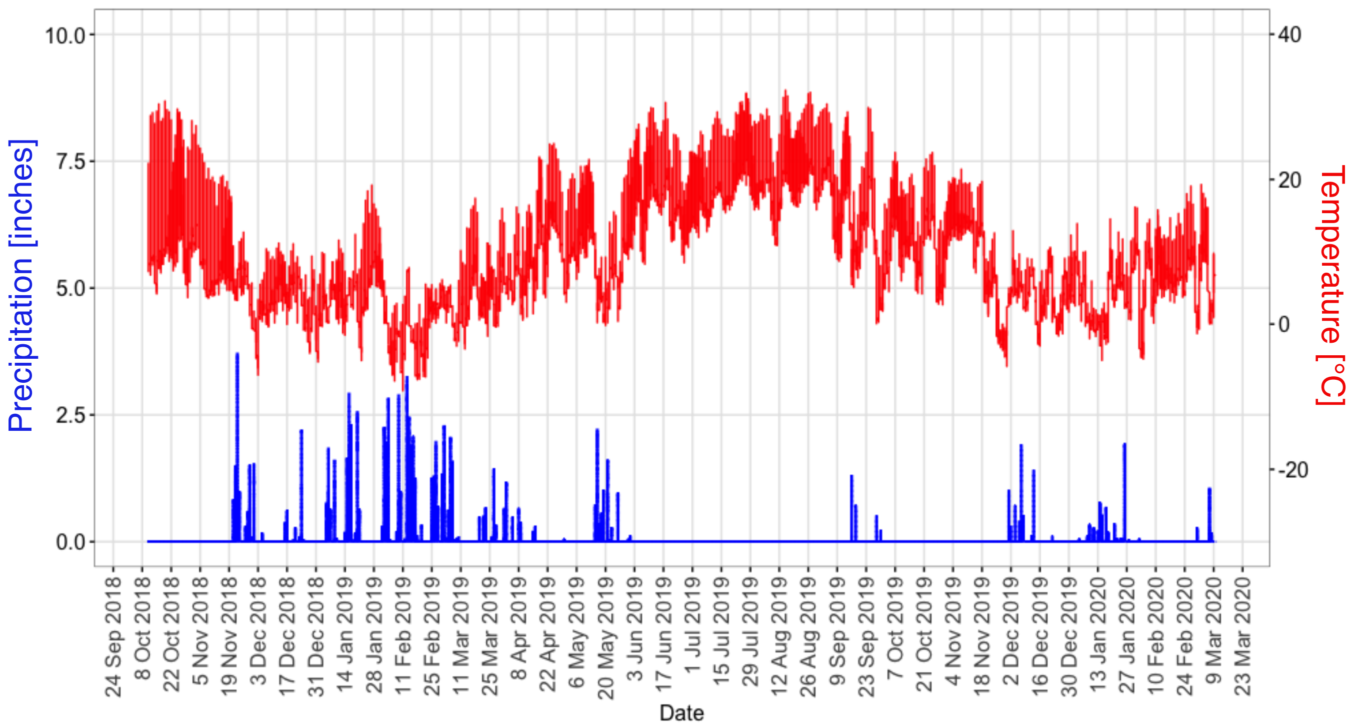


**Figure S8. Precipitation and temperature over time.** Precipitation and temperature data collected from the Blodgett forest weather station.
